# Sequence-specific minimizers via polar sets

**DOI:** 10.1101/2021.02.01.429246

**Authors:** Hongyu Zheng, Carl Kingsford, Guillaume Marçais

## Abstract

Minimizers are efficient methods to sample *k*-mers from genomic sequences that unconditionally preserve sufficiently long matches between sequences. Well-established methods to construct efficient minimizers focus on sampling fewer *k*-mers on a random sequence and use universal hitting sets (sets of *k*-mers that appear frequently enough) to upper bound the sketch size. In contrast, the problem of sequence-specific minimizers, which is to construct efficient minimizers to sample fewer *k*-mers on a specific sequence such as the reference genome, is less studied. Currently, the theoretical understanding of this problem is lacking, and existing methods do not specialize well to sketch specific sequences. We propose the concept of polar sets, complementary to the existing idea of universal hitting sets. Polar sets are *k*-mer sets that are spread out enough on the reference, and provably specialize well to specific sequences. Link energy measures how well spread out a polar set is, and with it, the sketch size can be bounded from above and below in a theoretically sound way. This allows for direct optimization of sketch size. We propose efficient heuristics to construct polar sets, and via experiments on the human reference genome, show their practical superiority in designing efficient sequence-specific minimizers. A reference implementation and code for analyses under an open-source license are at https://github.com/kingsford-group/polarset.

## 1 Introduction

The *minimizer* (Roberts *et al*., 2004a,b) methods, also known as *winnowing* (Schleimer *et al*., 2003), are methods to sample positions or *k*-mers (substrings of length *k*) from a long string. Thanks to its versatility, this method is used in many bioinformatics programs to reduce memory requirements and computational resources. Read mappers (Li and Birol, 2018; Jain *et al*., 2020b,a), *k*-mer counters (Erbert *et al*., 2017; Deorowicz *et al*., 2015), genome assemblers (Ye *et al*., 2012; Chikhi *et al*., 2015) and many more (see Marçais *et al*. (2019) for a review) use minimizers.

In most cases, sampling the smallest number of positions, as long as a string is roughly uniformly sampled, is desirable as it leads to sparser data structures or less computation as fewer *k*-mers need to be processed. Minimizers have such a guarantee of approximate uniform sampling: given the parameters *w* and *k*, it guarantees to select at least one *k*-mer in every window of *w* consecutive *k*-mers. It achieves this goal by selecting the smallest *k*-mer (the “minimizer”) in every *w*-long window, where smallest is defined by a choice of an order 𝒪 on the *k*-mers. Even though every minimizer scheme satisfies the constraint above, depending on the choice of the order 𝒪 the total number of selected *k*-mers may vary significantly.

Consequently, research on minimizers has focused on finding orders 𝒪 that obtain the lowest possible *density*, where the density is defined as the number of selected *k*-mers over the length of the sequence. In particular, most research concentrates on the average case: what is the lowest expected density given a long random input sequence? (Marçais *et al*., 2017, 2018; Ekim *et al*., 2020; Orenstein *et al*., 2016). In practice, many tools use a “random minimizer” where the order is defined by choosing at random a permutation of all the *k*-mers (e.g., by using a hash function on the *k*-mers). This choice has the advantage of being simple to implement and providing good performance on the average case.

Here we investigate a different setup that is common in bioinformatics applications. Instead of the average density over a random input we try to optimize the density for one particular string or sequence. When applying minimizers in computational genomics, in many scenarios the sequence is known well in advance and it does not change very often. For example, a read aligner may align reads repeatedly against the same reference genome (e.g., the human reference genome). In such cases, optimizing the density on this specific sequence is more meaningful than on a random sequence. Moreover, the human genome has markedly different properties than a random sequence and optimization for the average case may not carry over to this specific sequence. In the read aligner example, a minimizer with lower density leads to a smaller index to save on disk and fewer seeds to consider in the seed-and-extend alignment algorithm while preserving the same sensitivity thanks to the approximate uniform sampling property.

The idea of constructing sequence sketches tailored to a specific sequence has been explored before (Chikhi *et al*., 2015; DeBlasio *et al*., 2019; Jain *et al*., 2020b), but it remains less understood than the average case. Random sequences have nice properties that allow for simplified probabilistic analysis. Consequently, different analytic tools are needed to analyze sequence-specific minimizers. In fact, minimizers designed to have low density in the average case often offer only modest improvements on sequences of interest such as reference genomes (Zheng *et al*., 2020a).

The current theory for minimizers with low density in average is tightly linked to the theory of *universal hitting sets* (UHS) (Orenstein *et al*., 2016; Marçais *et al*., 2018; Kempa and Kociumaka, 2019). As the name suggests, a UHS is a set of *k*-mers that “hits” every *w*-long window of every possible sequence (hence the universality; it is an unavoidable set of *k*-mers). Universal hitting sets of small size generate minimizers with a provable upper-bound on their density. Universal hitting sets are less useful in the sequence-specific case as the requirement to hit every window of every sequence is too strong, and UHSs are too large to provide a meaningful upper-bound on the density in the sequence-specific case. New theoretical tools are needed to analyze the sequence-specific case.

Frequency-based orders are examples of sequence-specific minimizers (Chikhi *et al*., 2015; Jain *et al*., 2020b). In these constructions, *k*-mers that occur less frequently in the sequence compare less than *k*-mers that occur more frequently. The intuition is to select rare *k*-mers as they should be spread apart in the sequence, hence giving a sparse sampling. This intuition is only partially correct. First, there is no theoretical guarantee that a frequency-based order gives low density minimizers, and there are many theoretical counter-examples. Second, in practice, frequency-based orders often give minimizers with lower density, but not always. For example, Winnowmap (Jain *et al*., 2020b) uses a two-tier classification (very frequent vs. less frequent *k*-mers) as it performs better than an order strictly following frequency of occurrence.

Another approach to sequence-specific minimizers is to start from a UHS *U* and to remove as many *k*-mers from *U* as long as it still hits every *w*-long window of the sequence of interest (DeBlasio *et al*., 2019). Because this procedure starts with a UHS that is not related to the sequence, the amount of possible improvement in density is limited. Additionally, given the exponential growth in size of the UHS with *k*, current methods are computationally limited to *k* ≤ 15, which is limiting in many applications.

The construction proposed here takes a different approach and introduces *polar sets*. The polar sets concept can be seen as complementary to the universal hitting sets: while a UHS is a set of *k*-mers that intersects with every *w*-long window *at least* once, a polar set is a set of *k*-mers that intersect with any window *at most* once. The name “polar set” is an analogy to a set of polar opposite magnets that cannot be too close to one another. That is, our construction builds upon sets of *k*-mers that are sparse in the sequence of interest, and consequently the minimizers derived from these polar sets have provably tight bounds on their density.

Our main contribution is Theorem 1 that gives an upper bound and a lower bound on the density obtained by a minimizer created from a polar set. These bounds are expressed in term of the “total link energy” of the polar set on the given sequence. The link energy is a new concept that measures how well spread apart the elements of the polar sets are in the sequence: the higher the energy, the more spread apart the *k*-mers are. Then we show that the link energy is almost exactly the improvement in density one gains from using a minimizer created from the polar set compared to a random minimizer.

In the following sections we also show that the problem of finding a polar set with maximum total link energy is, unsurprisingly, NP-hard, and we describe a heuristic to create polar sets with high total link energy. Finally, we show that our implementation of this heuristic generates minimizers that have specific density on the human reference genome much lower than any other previous methods, and, for some parameter choices, relatively close to the theoretical minimum.

## 2 Methods

### 2.1 Overview

We set the stage by defining important terms and concepts, then giving an overview of the main results, which are then proved formally in the following sections. The sequence *S* is a string on the alphabet Σ of size *σ* = |Σ|. The parameters *k* and *w* define respectively the length of the *k*-mers and the window size. We assume that *S* is relatively long compared to these parameters: |*S*| ≫ *w* + *k*.

#### Definition 1

(Minimizer and Windows). *A minimizer is characterized by* (*w, k, 𝒪*) *where 𝒪 is a complete order of* Σ^*k*^. *A window is a sequence of length* (*w* + *k* − 1) *consisting of exactly w k-mers. Given a window as input, the minimizer outputs the location of the smallest k-mer according to 𝒪, breaking ties by preferring the leftmost k-mer*.

The minimizer (*w, k, 𝒪*) is applied to the sequence *S* by finding the position of the smallest *k*-mer in every window of *S*. Because two consecutive windows in *S* have a large overlap, the same *k*-mer is often selected in these two windows, hence the minimizer returns a sampling of positions in the sequence *S*. The *specific density* of the minimizer on *S* is defined as the number of selected positions over the length | *S* |.

The density is between 1*/w*, because at least one *k*-mer in every window must be picked, and 1, because it is a sampling of the positions of *S*. Therefore the goal is to find orders 𝒪 that have a density as close to 1*/w* as possible. A minimizer with density 1*/w* is a *perfect minimizer*. For simplicity, when stating the density of a minimizer we ignore any additive term that is *o*(1*/w*) (i.e., asymptotically negligible compared to 1*/w*).

A *random minimizer* is defined by choosing at random one of the permutations of all *k*-mers. The expected density of a random minimizer is 2*/*(*w* + 1) (Schleimer *et al*., 2003; Roberts *et al*., 2004b; Zheng *et al*., 2020a). Equivalently, the expected distance between adjacent selected *k*-mers is (*w* + 1)*/*2. The random minimizers will serve as a baseline to compare to.

**Defining orders**. For practical reasons, we define orders by defining a set *U* and considering orders that are *compatible* with *U*: an order 𝒪 is compatible with *U* if for 𝒪 every element of *U* compares less than any element not in *U*. That is, only the smallest elements for 𝒪 are specified (the elements of *U*) and a minimizer using an order compatible with *U* will preferentially select the elements of *U*. There exist many orders that are compatible with *U* as the relative order between the elements within *U* is not specified.

**Universal Hitting Sets**. A set *U* is a universal hitting if for every one of *σ*^*w*+*k−*1^ possible windows (recall *σ* is the size of the alphabet), it contains a *k*-mer from *U*. In the average case, minimizers compatible with *U* have densities upper bounded by |*U* |*/σ*^*k*^, because only *k*-mers from the universal hitting set can be selected. Supplementary Section S2 provides a more detailed discussion of why this bound provided by universal hitting sets does not always apply for sequence-specific minimizer analysis, and why universal hitting sets do not specialize well.

**Short sequences**. On a short random sequence (in a sense made precise by Lemma 1) most *k*-mers are unique (i.e., they occur only once in the sequence *S*). Therefore, it is likely that there is a set *U* of unique *k*-mers of *S* that are exactly *w* bases apart in *S*, and a minimizer compatible with *U* is perfect. Unfortunately most sequences of interest (e.g., reference genomes) are too long, too repetitive and in general do not satisfy the hypothesis of Lemma 1. For most sequences it is not possible to find a set of “perfect seeds” of *k*-mers spaced exactly *w* apart.

**Polar sets**. An *polar set* is a relaxed version of a perfect set: any pair of *k*-mers *m*_1_ and *m*_2_ from a polar set *A* are always more than *w/*2 bases apart in *S* (see the more general Definition 2). The intuition behind this definition is that for a minimizer compatible with *A*, any *k*-mer from *A* selected by the minimizer is at distance ≥ (*w* + 1)*/*2 from the previous and the next selected *k*-mer. Hence, *k*-mers selected from *A* are at least as sparse, and usually more sparse than *k*-mers selected using a random minimizer in expectation.

Section 2.2 gives a formal definition of the *link energy* of a polar set and Theorem 1 gives upper and lower bounds using this link energy for the density of a minimizer compatible with a polar set. This theorem shows that the link energy of the polar set *A* is a measure of how much reduction in density is obtained by using a minimizer compatible with *A* rather than a random minimizer. Hence, designing a polar set with high link energy is a method to find minimizers with provably low density.

Section 2.3 introduces *layered polar sets*, which are an extension to polar sets, and builds a heuristic method to create such sets.

### 2.2 Polar sets and link energy

#### 2.2.1 Key Definitions

We now define polar sets, the key component for our proposed methods.

##### Definition 2

(Polar set). *Given sequence S and parameters* (*w, k, s*) *with* 0 ≤ *s <* 1*/*2, *a* polar set *A of slackness s is a set of k-mers such that every two k-mers in A appears at least* (1 − *s*)*w bases apart in S*.

This can be viewed as a complementary idea to the universal hitting sets or a relaxed form of perfect sets. As discussed in the introduction, a universal hitting set requires the set to hit every *w* consecutive *k*-mers at least once, while a polar set with *s* = 0 requires the set to hit every *w* consecutive *k*-mers at most once. A set of perfect seeds, if it exists, is both a polar set with zero slackness and a universal hitting set. See Figure 1 for a more concrete example.

**Figure 1:**
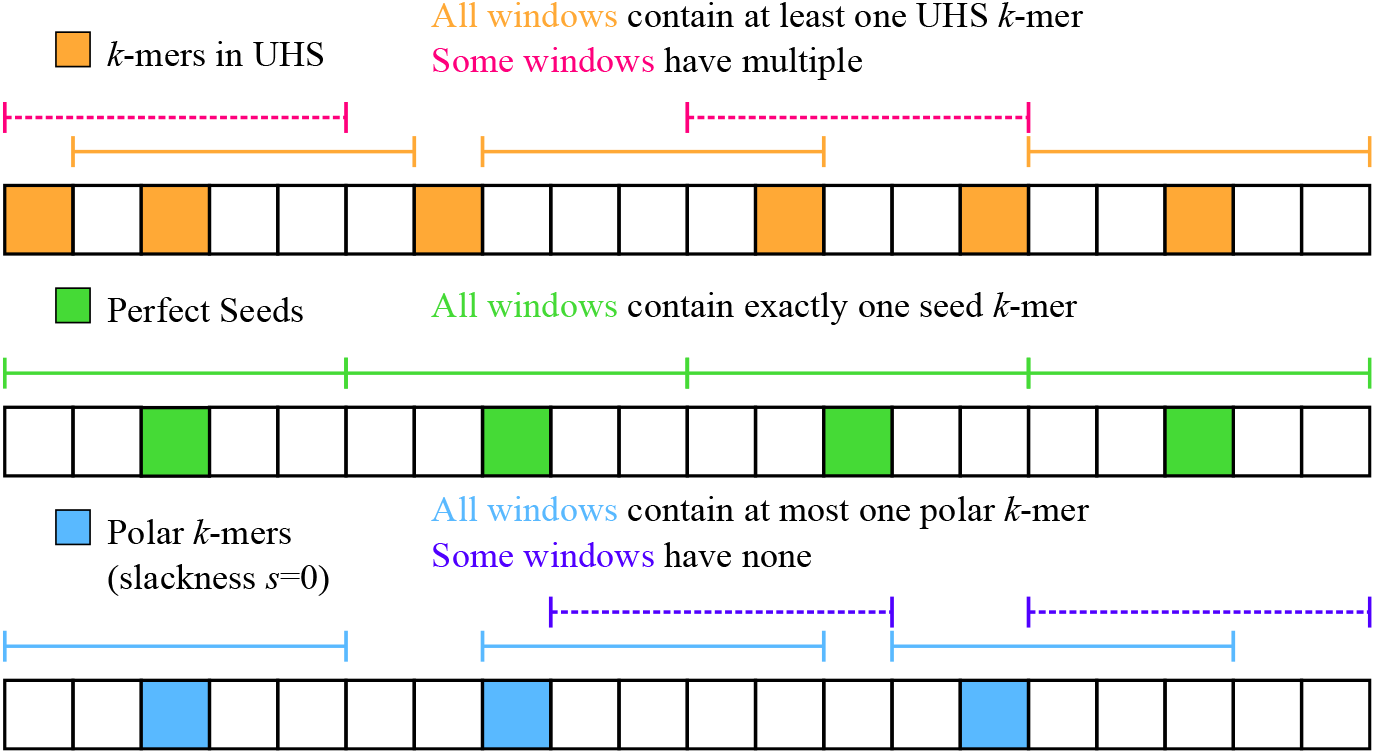
Comparing universal hitting sets, perfect seeds (compatible minimizers become perfect minimizers) and polar sets. Each block indicates a *k*-mer, and each segment indicates a window of length 5 (*w* = 5). To provide a better contrast with universal hitting sets, we show polar sets with slackness *s* = 0 (see Definition 2).

The condition *s <* 1*/*2 is critical for our analysis. Specifically, this condition is required to obtain a lower bound on the specific density of compatible minimizers, not just an upper bound.

##### Definition 3

(Link energy). *Given sequence S, parameters* (*w, k*) *and a polar set A, if two k-mers on S are l* ≤ *w bases apart and are both in A, the link energy of the pair is defined as* 2*l/*(*w* + 1) −1 ≥ 0. *The total link energy of A is the sum of link energy across all eligible pairs*.

Any two *k*-mers from *A* in *S* must be more than *w/*2 bases apart, so two *k*-mers cannot form a link if there is a third *k*-mer from *A* between them. With *s* = 0, the link energy is fixed to be 2*w/*(*w* + 1) −1 = 1− 2*/*(*w* + 1) ≈ 1 for each eligible pair, and the total link energy is approximately the number of pairs that form a link, which in turn is the number of *k*-mer pairs in the polar set that are exactly *w* bases away on *S*. In the following sections, we introduce and discuss the backbone of the polar set framework, which revolves around closer inspection of how a random minimizer works on a specific sequence, and drawing contrast between sequence-specific minimizers and non-sequence-specific minimizers. We use the term “non-sequence-specific minimizers” to refer to constructions of minimizer that does not specifically target a certain sequence, but rather aim to minimize density, the expected specific density on a random string.

#### 2.2.2 Perfect Minimizer for short sequences

A perfect minimizer is a minimizer that achieves density of exactly 1*/w*. While the only known examples of perfect minimizers are in the asymptotic case where *w* ≪ *k* (Marçais *et al*., 2018), perfect sequence-specific minimizers exist with high probability for short sequences.

##### Lemma 1.

*If* 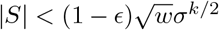, *with at least ϵ probability a random sequence of length* |*S* | *has a perfect minimizer*.

*Proof*. The optimal minimizer is constructed with fixed interval sampling. More specifically, we take every *w k*-mer in *S* and denote the resulting *k*-mer set *U*, then construct a minimizer compatible with *U*. The resulting minimizer is perfect if and only if the *k*-mers in *U* only appear in the selected locations. There are |*S* |*/w* selected locations and (1 −1*/w*) |*S* | locations not selected, and for each pair of selected and not selected locations, the *k*-mer at these two locations are identical with probability *σ*^*−k*^ (see Supplementary Section S1). By union bound, the probability that the sequence violates the polar set condition is at most |*S*|2*σ*^*−k*^*/w <* (1 − ϵ)^2^, and the sequence has a perfect minimizer with probability at least 1 −(1 − ϵ)^2^ *> ϵ*.

#### 2.2.3 Context Energy and Energy Savers

*Contexts* provide an alternative way to measure the density of a minimizer (Zheng *et al*., 2020a). These play a central role on the analysis of polar sets.

##### Definition 4

(Charged Contexts). *A* context *of S is a substring of length* (*w* + *k*), *or equivalently* (*w* + 1) *consecutive k-mers, or equivalently* 2 *consecutive windows*.

*A context is* charged *if the minimizer selects a different k-mer in the first window than in the second window*.

See top left of Figure 2 for examples of charged contexts. Intuitively, a charged context corresponds to the event that a new *k*-mer is picked, and counting picked *k*-mers is equivalent to counting charged contexts.

**Figure 2:**
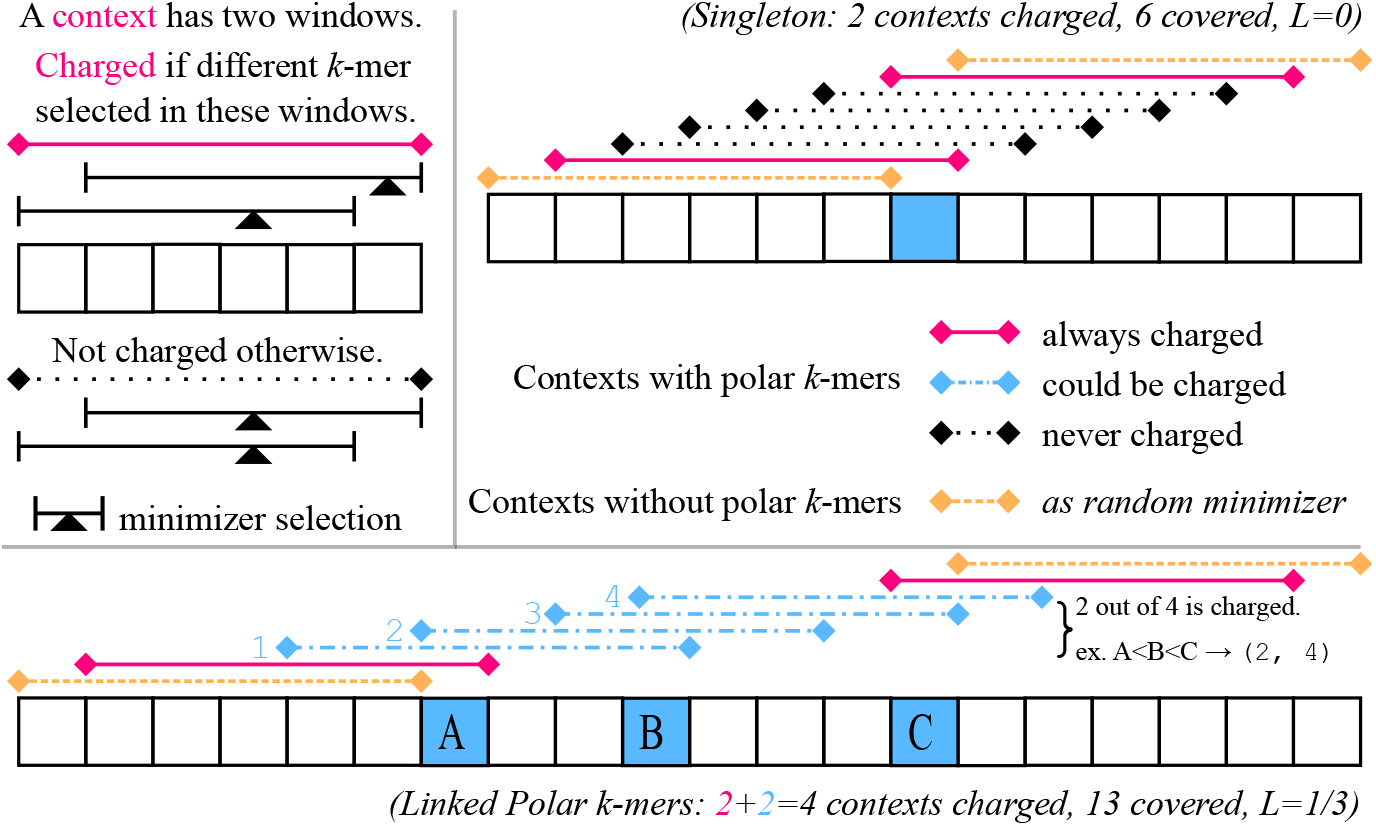
Examples for our argument for the polar set density bound with *w* = 5. Left top: Legend for a context, and when it is charged. Right top: Case for a singleton polar *k*-mer without links. In this case, *L*(*S, A*) = 0. Bottom: Case for three linked polar *k*-mers. Whatever the ordering between the three polar *k*-mers, two of the four contexts marked in blue will be charged. The link energy *L*(*S, A*) = 1*/*3: *A* and *B* are *l* = 3 bases away with no energy, *B* and *C* are *l* = 4 bases away with energy 2*l/*(*w* + 1) − 1 = 1*/*3.

##### Lemma 2

(Specific Density by Charged Contexts). *For a given sequence S and a minimizer, the number of selected locations by the minimizer equals the number of charged contexts plus* 1.

Given a context *c*, define *E*(*c*) as the probability that *c* will be charged with a random minimizer (one with a random ordering of *k*-mers), which we call the *energy* of *c*.

##### Lemma 3.

*The expected number of picked k-mers in S under a random minimizer is* 1 + *E*_0_(*S*), *where E*_0_(*S*) = *c E*(*c*) *is called the initial energy of S and the summation is over every context of S*.

This is proved by combining the linearity of expectation and Lemma 2. This implies that the total energy of a sequence is directly related to the specific density of random minimizers, which is number of picked locations in *S* divided by number of *k*-mers in *S. E*(*c*) admits a simple formula:

##### Lemma 4.

*E*(*c*) = 2*/u*(*c*) *if the last k-mer in the context is unique*, 1*/u*(*c*) *otherwise, where u*(*c*) *denotes the number of unique k-mers in c*.

*Proof*. Consider an imaginary minimizer with *w*^*′*^ = *w* +1 and identical *k*. The context of a (*w, k*) − minimizer is a window of the imaginary minimizer, and it is charged if and only if the imaginary minimizer picks either the first or the last *k*-mer. If the imaginary minimizer does not pick either end, the two constituent windows of the context share the same minimal *k*-mer, and the context is not charged.

With a random minimizer, the probability that the first *k*-mer is picked in the imaginary window is 1*/u*(*c*). The probability that the last *k*-mer is picked is 1*/u*(*c*) if the last *k*-mer is unique, 0 otherwise, because the minimizer break ties by preferring leftmost *k*-mer. The two events are mutually exclusive, so *E*(*c*) is the sum of these two terms.

If all *k*-mers in a context are unique, *E*(*c*) = 2*/*(*w* + 1) is guaranteed, which we call the baseline. If this holds for all windows, a random minimizer will have specific density of 2*/*(*w* + 1), similar to applying a random minimizer to a random sequence. As lower *u*(*c*) only increases *E*(*c*), *E*(*c*) *<* 2*/*(*w* + 1) only if the last *k*-mer in *c* is not unique and there are over (*w* + 1)*/*2 unique *k*-mers in the context.

**Definition 5**. *A context c is called an* energy saver *if E*(*c*) *<* 2*/*(*w* + 1), *and its energy deficit is defined as* 2*/*(*w* + 1) − *E*(*c*). *The energy deficit of S, denoted D*(*S*), *is the total energy deficit across all energy savers: D*(*S*) = ? _*c*_ max(0, 2*/*(*w* + 1) − *E*(*c*)).

In general, the value of *D*(*S*) is very small due to the fact that energy saver contexts (those with *E*(*c*) *<* 2*/*(*w* + 1)) are rare.

##### Lemma 5.

*For a random context, the probability that it is an energy saver is at most wσ*^*−k*^.

*Proof*. We bound the probability that the last *k*-mer in a context is not unique. The probability that the last *k*-mer equals a specific *k*-mer in another location is *σ*^*−k*^ (see Supplementary Section S1). Applying union bound over *w* other *k*-mers (as each context has (*w* + 1) *k*-mers) we get the desired result.

There are examples of sequences where energy saver contexts are abundant. An extreme scenario is when the sequence *S* is has a period of *w*, and has *w* distinct *k*-mers. In this case, all contexts become energy saver contexts. These scenarios are rare in practice.

Similarly, we can define energy spenders and energy surplus as follows:

##### Definition 6.

*A context c is called an* energy spender *if E*(*c*) *>* 2*/*(*w* + 1), *and its surplus is defined as E*(*c*) − 2*/*(*w* + 1). *The energy surplus of S, denoted X*(*S*), *is the total energy surplus across all energy spenders: X*(*S*) =∑ _*c*_ max(0, *E*(*c*) − 2*/*(*w* + 1)).

Contexts with energy surpluses are more common than energy savers, but still fairly rare in a random sequence with suitable choice of *w* and *k*:

##### Lemma 6.

*For a random context, the probability that it is an energy spender is at most w*(*w* + 1)*σ*^*−k*^*/*2.

*Proof*. A context becomes an energy spender if the last *k*-mer is unique, and some *k*-mers appears twice. We bound the probability that some *k*-mers in the context appear twice. Following previous arguments, any two *k*-mers in a given context are identical to each other with probability *σ*^*−k*^, and we apply a union bound of size *w*(*w* + 1)*/*2 (enumerating over pairs of *k*-mers) to obtain the desired result.

#### 2.2.4 Density Bounds with Polar Sets

With the proper tools, we now state the main theorem of the Polar Sets.

##### Theorem 1.

*Given a sequence S and a polar set A on S, let E*_0_(*S*) *be the initial energy of S, D*(*S*) *be the total energy deficit, X*(*S*) *be the total energy surplus, and L*(*S, A*) *be the total link energy from the polar set. The number of selected k-mers over S for a random minimizer compatible with A is at most* 1 + *E*_0_(*S*) + *D*(*S*) − *L*(*S, A*), *and at least* 1 + *E*_0_(*S*) − *X*(*S*) − *L*(*S, A*).

*Proof*. We first prove the upper bound part. We start by elevating the energy of every energy saver context to the baseline 2*/*(*w* + 1). By definition, this increases the total energy of *S* by *D*(*S*), so number of selected *k*-mers is now upper bounded by 1 + *E*_0_(*S*) + *D*(*S*). Formally, *E*(*x*) ≤ 1 + *E*_0_(*S*) + *D*(*S*).

Consider the minimizer compatible with *A*, with arbitrary ordering within *A* and random ordering outside *A*. We can still calculate the expected number of selected *k*-mers by summing up the probability of every context being charged, which we denote *E*_*A*_(*x*). Our goal for the rest of this proof is to show ∑ (*E*(*x*) − *E*_*A*_(*x*)) ≥ *L*(*S, A*).

If a context does not contain a *k*-mer from *A, E*(*x*) = *E*_*A*_(*x*). Thus, we only need to consider the contexts that contain at least a *k*-mer from *A*, which are split into continious segments by the set of contexts not containing *k*-mers from *A*.

If a segment only contains one *k*-mer from *A*, there are exactly (*w* + 1) contexts in this segment (see Figure 2, upper right for an example). As each context now has energy at least 2*/*(*w* + 1), the total energy from (*w* + 1) contexts is at least 2. However, *E*_*A*_(*x*) across these contexts is exactly 2, as exactly two contexts will be always charged (the first and last in the segment), and every other context will never be charged. This means such segments can be ignored in the upper bound analysis, as it can only decrease the total energy: (*E*(*x*) −*E*_*A*_(*x*)) ≥ 0.

We now focus on the segments with more than one *k*-mer from *A* (see Figure 2, bottom for an example). Let *n* be the number of contexts from this segment, we have *E*(*x*) ≥ 2*n/*(*w* + 1) because we have assumed *E*(*x*) ≥ 2*/*(*w* + 1) for every context. We next count the number of charged contexts, which is *E*_*A*_(*x*). Because every two *k*-mers in *A* are more than *w/*2 bases apart, for every *k*-mer *m* in *A*, there is a window such that *m* is the only polar *k*-mer, and will be selected. Recall a context is charged if and only if a new *k*-mer is selected in its latter window. Given that each of the *d k*-mers in *A* is selected, this corresponds to *d* charged contexts in the segment. However, there is an extra charged context: The last context *c* in the segment, of which the last *k*-mer in *A* is the first *k*-mer of *c*. In the second window of *c*, a new *k*-mer outside *A* will be selected because being at the end of the segment, there are no more *k*-mers in *A* to choose from. Let *d* be the number of *k*-mers in *A* in this segment, we conclude that *E*_*A*_(*x*) = *d* + 1 regardless of the ordering, and (*E*(*x*) − *E*_*A*_(*x*)) ≥ 2*n/*(*w* + 1) − *d* − 1.

Next, we calculate the total link energy. Recall the link energy is defined as 2*l/*(*w* + 1) − 1 for two polar *k*-mers *l* ≤ *w* bases away. The total link energy is thus 2(*l*)*/*(*w* + 1) − (∑1), summed across all *k*-mer links. The latter term is simply counting number of links, which resolves to *d* − 1. The earlier term is summing up distance between adjacent *k*-mers in *A*. As this is a segment where every two adjacent *k*-mers have a link, *l* is the distance from the first *k*-mer in *A* to the last *k*-mer in *A*. There are *n* contexts in the segment. The first *k*-mer in *A* is the last *k*-mer in the first context, and the last *k*-mer in *A* is the first *k*-mer in the last context. The first context and the last context are *n* − 1 bases away. The first *k*-mer and the last *k*-mer in a context are *w* bases away. Thus, we have *∑ l* = *n* − 1 − *w*.

Finally, we have the following, using *S*^*l*^ to denote the sequence that contains the contexts in a segment:

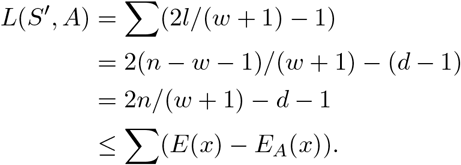

The summation is over each context in the segment. This inequality holds for every segment, and thus we have:

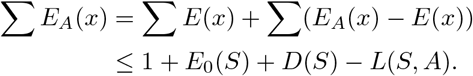

The lower bound uses a symmetric argument as we first upper bound the energy of each context by the baseline 2*/*(*w* + 1). This decreases total energy by *X*(*S*) and total expected number of selected *k*-mers is lower bounded by 1 + *E*_0_(*S*) − *X*(*S*). The identical argument (with signs flipped) will lead to the final lower bound *E*_*A*_(*x*) ≥ 1 + *E*_0_(*S*) − *X*(*S*) − *L*(*S, A*).

As 1 + *E*_0_(*S*) is the expected number of selected *k*-mers with a completely random minimizer, we can provably outperform random minimizers if *L*(*S, A*) *> D*(*S*). For a ballpark estimate, we assume *S* is a random sequence, and assume the slackness parameter *s* = 0 in construction of the polar set. In this setup, each link has exactly 1 − 2*/*(*w* + 1) ≈ 1 energy. As seen in Lemma 5, a context is an energy saver with probability *wσ*^*−k*^, and its deficit is at most 2*/*(*w* + 1) −1*/w* ≈ 1*/w*, meaning *D*(*S*) ≈ *σ*^*−k*^ |*S*|. This further means we need the number of links to be at least *σ*^*−k*^ *S* to provably beat a random minimizer. On the other hand, ignoring the effect of *D*(*S*), in order to beat the specific density of a random minimizer by ϵ*/*(*w* + 1), total link energy of ϵ |*S* | */*(*w* + 1) is needed. Assuming no slackness, this means the number of links need to be at least ϵ |*S* |*/*(*w* − 1). Intuitively, ϵ portion of the sequence needs to be covered by links between close enough *k*-mers in polar set.

A proper polar set requires *s >* 1*/*2 for the main theorem to hold. When *s* ≤ 1*/*2, only the upper bound part of the theorem holds with an alternative definition of link energy. We will discuss the alternative definition in Section 2.3.4, and further discuss generalization of polar sets in Supplementary Section S2.4.

#### 2.2.5 Hardness of Optimizing Polar Sets

The link energy formulation of polar sets allows us to cast the problem in graph theoretical framework. Consider an undirected, weighted graph where every unique *k*-mer is a vertex. An edge connects two *k*-mers with the following: If these two *k*-mers ever appear within fewer than (1− *s*)*w* bases of each other in *S*, the weight is −∞. Otherwise, the weight of this edge is the total link energy by selecting only these two *k*-mers, which might establish several links given each *k*-mer may appear in *S* multiple times. There can also be self-loops with weights, given a *k*-mer may appear close to itself on the reference sequence. The problem of finding optimal polar sets becomes the problem of finding an induced subgraph with maximum weight.

The general maximum induced subgraph problem is well known to be NP-hard via reduction from max-clique. In Supplementary Section S3, we provide an explicit proof that shows optimization of polar sets, even with an alphabet of three, is NP-Hard.

### 2.3 Constructing Polar Sets

In this section, we propose a practical extension to polar sets, and formally introduce our heuristics.

#### 2.3.1 Layered Polar Sets

Assume we have already constructed a polar set *A* that covers some segments of the reference sequence. Here, covered means that every window contains a *k*-mer from the polar set, or equivalently, *A* acts as a universal hitting set on these segments.

Now, to cover the rest of the reference, we shall extend *A* so more *k*-mers become polar *k*-mers. It is natural to consider generating a polar set over the uncovered portion of the reference sequence, then merge this set with *A*. This however leads to problems. Let *A*^*′*^ be a polar set over the uncovered portion of the reference sequence. *A* ∪ *A*^*′*^ might not always be a valid polar set, because a *k*-mer *m*^*′*^ ∈ *A*^*′*^ may appear in the already-covered part of the reference sequence, and appear close to another *k*-mer *m* ∈ *A*, thus violating the polar set condition for *A* ∪ *A*^*l*^.

On the other hand, the reason we set up the constraint for polar sets is to ensure that *k*-mers in the polar set will always be selected by any compatible minimizer. In other words, we want to ensure we know exactly the set of *k*-mers that will be selected. The issue was that *m*^*′*^ ∈ *A*^*′*^ might not always be selected by a compatible minimizer. However, from the perspective of constructing efficient minimizers, we do not need *m*^*′*^ to be selected everywhere, as in some places the reference sequence is already covered with *k*-mers in *A*. By forcing *m < m*^*′*^ for any *m* ∈ *A*, we ensure that *m*^*′*^ will only be selected outside the segments covered by *A*.

Applying this argument to all *k*-mers in *A*^*′*^, we can essentially ignore the sequence segments already covered by *A* when constructing *A*^*′*^, as long as the ordering is satisfied. This gives a way to progressively construct the layers of polar sets: at each layer we only need to consider regions of the reference sequence that are not yet covered by previous layers. Formally:

##### Definition 7.

*A layered polar set is a list of sets of k-mers* {*A*_*i*_}, *for* 1 ≤ *i* ≤ *m. With slackness s, the layered polar set condition is satisfied if for any k-mer in A*_*j*_, *for each of its appearance at location t in the reference sequence, either of the following holds:*

*It is at least* (1 − *s*)*w bases apart from any k-mer in* {*A*_1_, *A*_2_, ⃛, *A*_*j*_}.

- *It is covered: There are two k-mers in* {*A*_1_, *A*_2_, ⃛, *A*_*j−*1_} *(importantly does not include A*_*j*_*), appearing at location l and h, satisfying l < t < h and h* − *l* ≤ *w*.

Similarly, a compatible order for {*A*_*i*_} is an order that places all *k*-mers from *A*_1_ first in arbitrary order, then those in *A*_2_, …, then those in *A*_*m*_ and finally those not in any of {*A*_*i*_} in a random order. The link energy *L*({*A*_*i*_}, *S*) is similarly defined over the pairs of close *k*-mer appearances that are not covered. More formally:

##### Definition 8.

*For a layered polar set, if two k-mers in the layered polar sets, not necessarily from the same layer, appear l* ≤ *w bases apart in S, and neither are covered, the link energy between them is* 2*l/*(*w* +1)−1 *>* 0. *L*({*A*_*i*_}, *S*) *is the total link energy across all pairs*.

These definitions of layered polar sets and link energy have two important properties. First the link energy is non-decreasing as more layers are added to the set. And, second, an almost identical argument proves the same bounds for layered polar sets as for polar sets in Theorem 1. See Figure 3 for a concrete example of layered polar sets.

**Figure 3:**
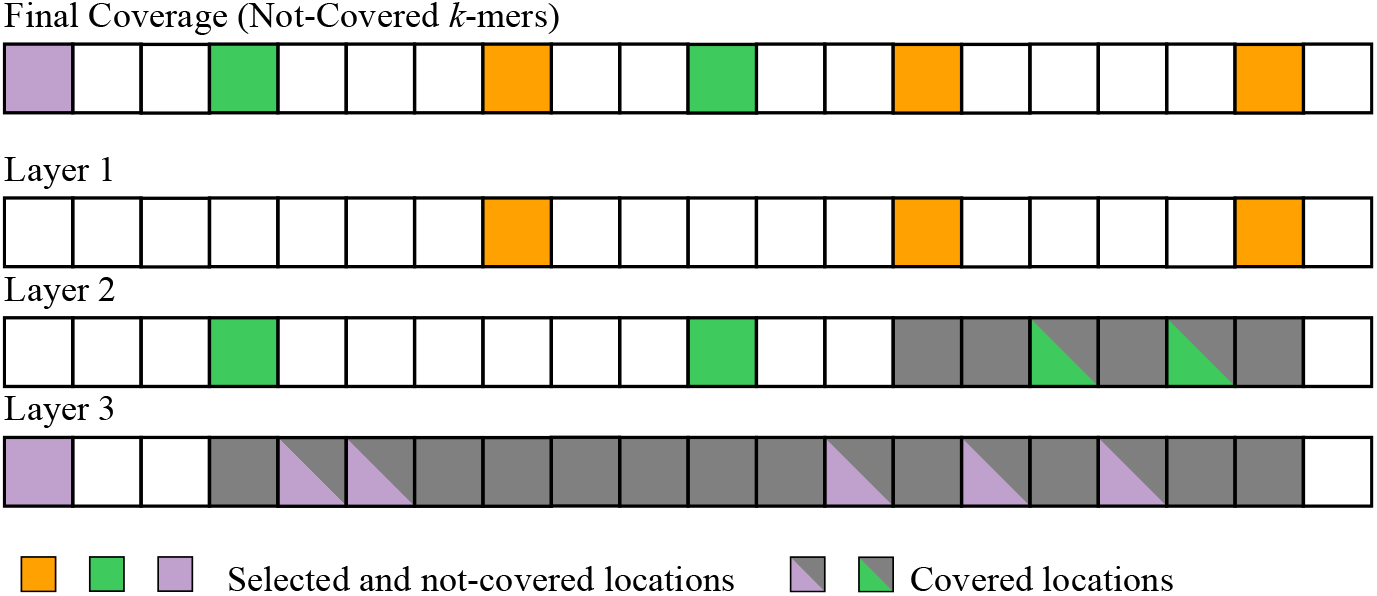
Examples of layered polar sets, with three layers. Without layered polar sets, the *k*-mers from layer 2 and 3 could not be selected as in the polar set because of self-collision. The whole sequence is covered in this case (every window contains a polar *k*-mer from one layer). Layer 1 is the one with highest priority and our layered heuristics construct it first.

#### 2.3.2 Polar Set Heuristic

We consider a simple heuristic to generate a polar set. The core idea is to select as many *k*-mers as possible from the set of *k*-mers that appear exactly *w* bases away from each other. We cannot select all of them as it may violate the polar set condition due to some *k*-mers appearing multiple times. Because reference sequences are long strings (in the range of billions of bases for mammalian genomes), we consider algorithms that scale well with the length of the reference sequence, preferably close to linear.

Fix an offset *o* ∈ [0, *w* − 1], we start by listing all locations *t* such that *t* = *o* mod *w* in the reference sequence *S*. We then randomly shuffle the locations, and for each location *t* in this random order, add the *k*-mer at location *t* to the polar set. When we add a *k*-mer *m* to the polar set, we also locate and remove all *k*-mers in the polar set that appear fewer than (1 − *s*)*w* bases away from *m*. Additionally, if a *k*-mer appears multiple times in the list, it is considered only once at the first encounter. This is to prioritize *k*-mers that appear less often; Frequent *k*-mers are expected to be processed early given their multiple occurrences, and are more likely to be absent in the final polar set as they have more chances to be removed due to conflicts. Jain *et al*. (2020b) has explored a similar idea in building tiered random minimizers using a biased hash function.

Our algorithm also has a variant, which we call “monotonic”. In this variant, we require that adding a new *k*-mer *m* and removing the *k*-mers conflicting with *m* actually increases the link energy. Otherwise, the *k*-mer is skipped and no conflicting *k*-mers are removed. This variant is slower but results in more efficient polar sets.

We filter *k*-mers before they are considered for addition to polar sets. *k*-mers that collides with itself (appears fewer than (1− *s*)*w* bases away from its own copy) cannot be in the polar set. We also filter out *k*-mers by their frequency in the reference sequence (see Section 2.3.3 for the threshold value).

##### Algorithm 1 Pseudocode for Polar Set Heuristics

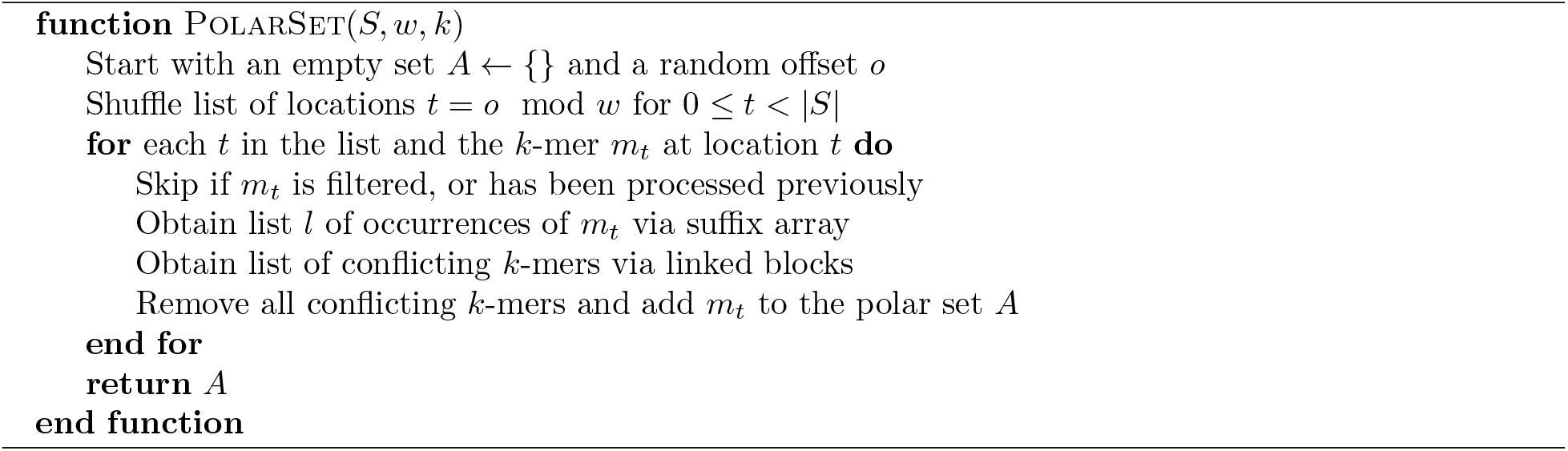

Algorithm 1 shows the pseudocode for the non-monotonic variant of the heuristic. The monotonic variant is similar. We describe the data structures in Section 2.3.4, and analyze the time complexity in Section 2.3.5.

#### 2.3.3 Layered Heuristics and Hyperparameters

We construct layered polar sets with a similar algorithm. The properties of layered polar sets guarantee that new layers cannot decrease the final link energy of the polar set.

We rerun the polar set heuristic multiple times, each time with a new random offset *o*. Each round is run with the current layers of polar sets, and the resulting polar set is added as a new layer. The algorithm for each layer is mostly identical to the single-layer version, with a few changes.

- When processing a *k*-mer, we skip all of its occurrences that are covered by existing layers of the polar set.
- We skip *k*-mers at non-covered locations *t* that is fewer than (1 − *s*)*w* bases away from a *k*-mer in a previous layer. These *k*-mers cannot be in the layer without violating the layered polar set condition.
- At the end of each round, we remove all *k*-mers selected in the current layer that do not form a link with any *k*-mers.

We also gradually increase the threshold of *k*-mer frequency at each round to prioritize low frequency *k*-mers. In our experiments, we use a total of 7 rounds, with last two rounds being monotonic. The frequency threshold is set at the value to include 85% of locations of the reference in the first round, gradually increasing to 95% in the last round.

The slackness *s* is also a tunable parameter, which determines when a pair of *k*-mers is considered in collision. Lower value of *s* ensures the distance between adjacent polar *k*-mers are large and have higher link energy for every pair of linked *k*-mers, but results in smaller number of *k*-mers selected, implying fewer links. Higher value of *s* means larger polar sets covering more of the reference sequence and more links formed, but adjacent polar *k*-mers may be closer to each other resulting in lower energy per link.

In our experiments, we use a fixed slackness *s* = 0.4 after parameter search. This results in approximately 20% less efficient links (average link energy compared to theoretical maximum), but higher total link energy due to inclusion of more links. A more thorough parameter tuning might suggest a gradually increasing value of *s* between rounds.

#### 2.3.4 Supporting Data Structures

Our heuristics require some data structures to operate efficiently both in theory and in practice.

**Suffix Array**. In order to quickly index *k*-mers and obtain the list of occurrences of a *k*-mer, we precompute the suffix array, the inverse suffix array and the heights table (also known as the LCP array) of the reference sequence. All can be computed in linear time. This allows us to find the list of *T* locations that share the same *k*-mer as location *t*, in *O*(*T*) time.

**Linked Blocks**. The layered polar set property ensures that in any stretch of *w/*2 bases, at most one *k*-mer at one location is selected into any layer of the layered polar sets, excluding covered locations. We use a data structure called linked blocks to represent the set of these selected locations of *k*-mers. Let *h* = ⌊*w/*2 ⌋, we divide the locations in the reference sequence into *h*-long blocks, and use an array of length |*S* |*/h* to represent these blocks. Each value in the array *C*[*b*] is either 1, meaning there are no selected location within this block spanning location [*bh*, (*b* + 1)*h*), or a nonnegative integer *j*, indicating that the *k*-mer at location *bh* + *j* is selected. With linked blocks we can do the following operation quickly:

##### Definition 9.

*PeekL(x) returns the closest selected location to the left of x, up to w bases*.

This is because we only need to query up to three blocks. Adding a location and removing a location also only involves a single block. Similarly we can define PeekR(x). With this data structure, we can implement many critical operations in the aforementioned heuristics. The step of filtering *k*-mers, more specifically determining whether a *k*-mer collides with itself, is done using this data structure, in similar fashion to bucket sorting. By maintaining two linked blocks, one for the current layer and one for all previous layers, we can determine whether a location is covered by the previous layers, and list collisions on the current layer.

**Calculating Link Energy**. In the monotonic variant of our heuristics, we need to calculate the total link energy before and after adding a *k*-mer. In our implementation, we update the link energy of the polar set as we add and remove locations to the linked blocks, using the following alternative formula for link energy:

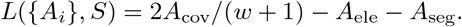

Here, *A*_cov_ is the number of contexts that contain a *k*-mer from the polar set, *A*_ele_ is the number of non-covered location of selected *k*-mers, and *A*_seg_ is the number of continuous segments of windows that contain a *k*-mer from the polar set. When adding and removing a location to the linked blocks, the changes to these three values are calculated using linked block primitives in constant time, so we can update the link energy in constant overhead. As a sanity check, we see that when adding an isolated *k*-mer, *A*_cov_ increases by (*w* + 1) and the other two values increase by 1, resulting in a net link energy gain of zero, consistent with the original definition. We can also compute the link energy of the polar *k*-mers in bottom part of Figure 3 using this formula, where *A*_cov_ = 13, *A*_ele_ = 3 and *A*_seg_ = 1, resulting in the total link energy of 1*/*3.

#### 2.3.5 Time Complexity Analysis

We now analyze the time complexity of the layered polar sets heuristic, assuming no monotonic rounds for now. Let *n* be the length of the reference sequence, and assume a constant-sized alphabet. We assume a word of constant size can hold an integer in [0, *n*], and that accessing an element in an array of length *n* takes constant time. These conditions hold for genomes and 64-bit machines. This means the primitive operations on linked blocks take constant time, and operations involving the suffix array also take constant time.

Consider a worst case scenario: By iterating *k*-mers that appear exactly *w* bases away from each other, we iterated over all *k*-mers in the reference sequence. Assume a *k*-mer *m* occurs *T* times in the reference sequence. In filtering phase, we first fetch the list of *T* locations in *O*(*T*) time using the suffix array, and we want to determine if there are two elements whose difference is less than (1− *s*)*w*. This can be done using the linked blocks in *O*(*T*) time. In the case of layered polar sets, we also want to determine if each of the locations is covered by previous layers, and if it is fewer than (1 − *s*)*w* bases away from a location in a previous layer. As we use one linked block for all previous layers, this can be done in *O*(*T*) time. The filtering phase thus finishes in *O*(*T*) time.

The main algorithm is split into three parts: detecting *k*-mers that are close to *m* in the reference sequence, removing those *k*-mers from the polar set, and adding *m* to the polar set. Detecting and listing *k*-mers that are close to *m* takes *O*(*T*) time, as each location reports only four collisions at most, two to the left and two to the right. Removing a *k*-mer that occurs *T* ^*l*^ times takes *O*(*T* ^*l*^) time, but since each *k*-mer is only added and removed once in one round, this amortizes to *O*(*T*) time. Adding *m* to the polar set also takes *O*(*T*) time. The singleton detection step (removing *k*-mers forming no links) also takes *O*(*T*) time for checking if *m* is a singleton.

As each *k*-mer is only visited once in the main algorithm, and in the worst case scenario every *k*-mer in *S* is visited, we conclude that the layered polar set heuristics runs in *O*(*T*) = *O*(*n*) time for each layer, and as a special case the (non-layered) polar set heuristics runs in *O*(*n*). The monotonic variant of the heuristic can in theory run in *O*(*n*^2^) time, but it is not significantly slower in practice.

## 3 Results

All the experiments are run using the human reference genome hg38. To facilitate the performance comparison across a range of parameter values of *w* and *k*, we report the *density factor* (Marçais *et al*., 2017) instead of the density. The density factor is the density multiplied by (*w* + 1). Regardless of the value of *w*, the random minimizer has an expected density factor of 2 and a perfect minimizer has a density factor of ≈ 1.

### 3.1 Energy Deficit and Energy Surplus

First, we calculate the average energy deficit *X*(*S*)*/* |*S*| and average energy surplus *D*(*S*)*/* |*S*|. The results are in Figure 4A.

**Figure 4:**
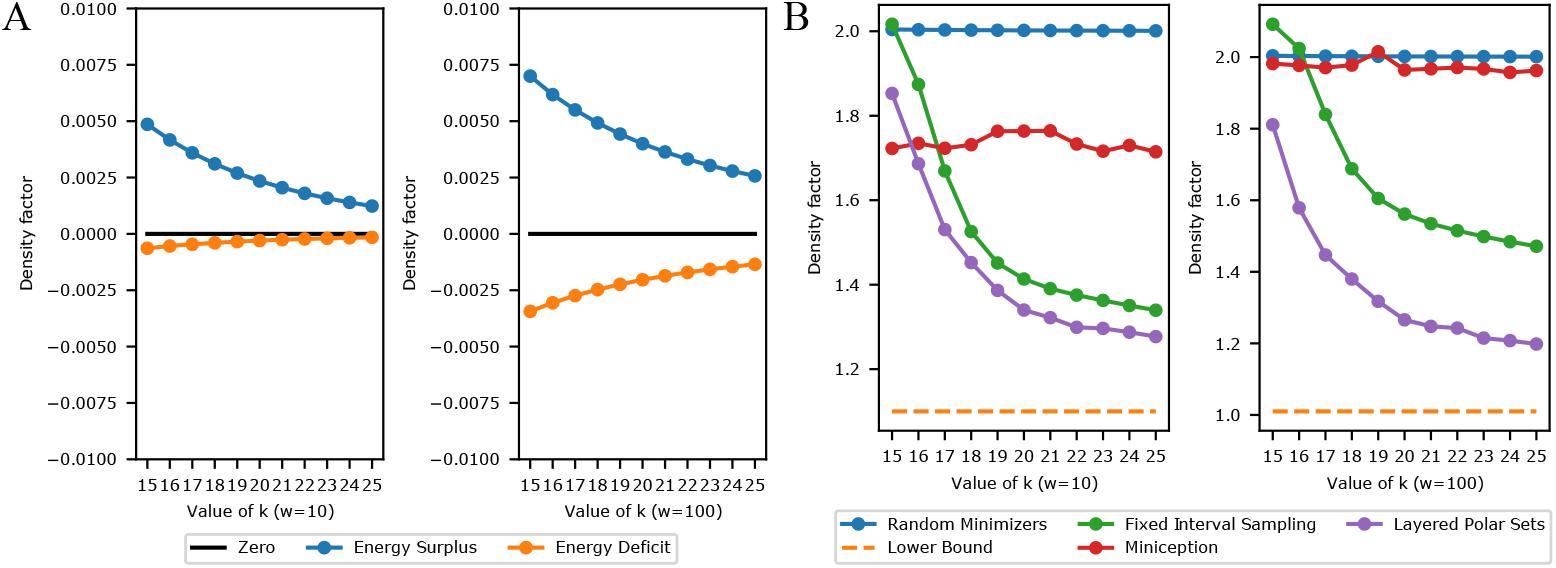
Left: Energy surplus and deficit for short (*w* = 10) and for long (*w* = 100) windows, computed on the human reference sequence hg38. The difference between the two lines is the difference between the upper and lower bound of Theorem 1. It is very small and the bounds are very good estimates in practice. Right: Density factor for the proposed methods, for short and long windows, computed on hg38. The bottom orange dashed line is the theoretical minimum density (perfect minimizers).

**Figure 5:**
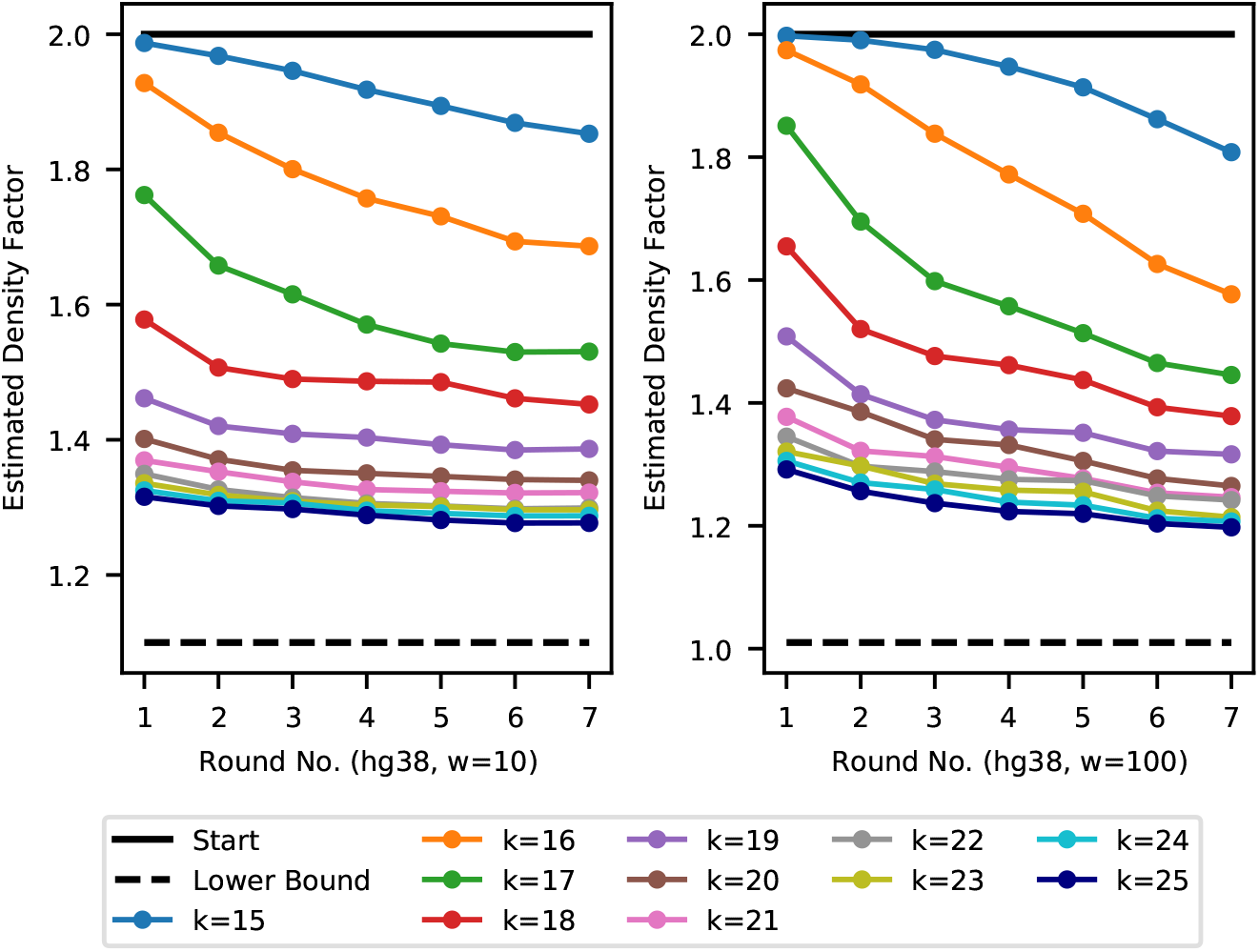
Density factor of layered anchor sets after each round of the optimization, corresponding to the experiments shown in Figure 4B.

**Figure 6:**
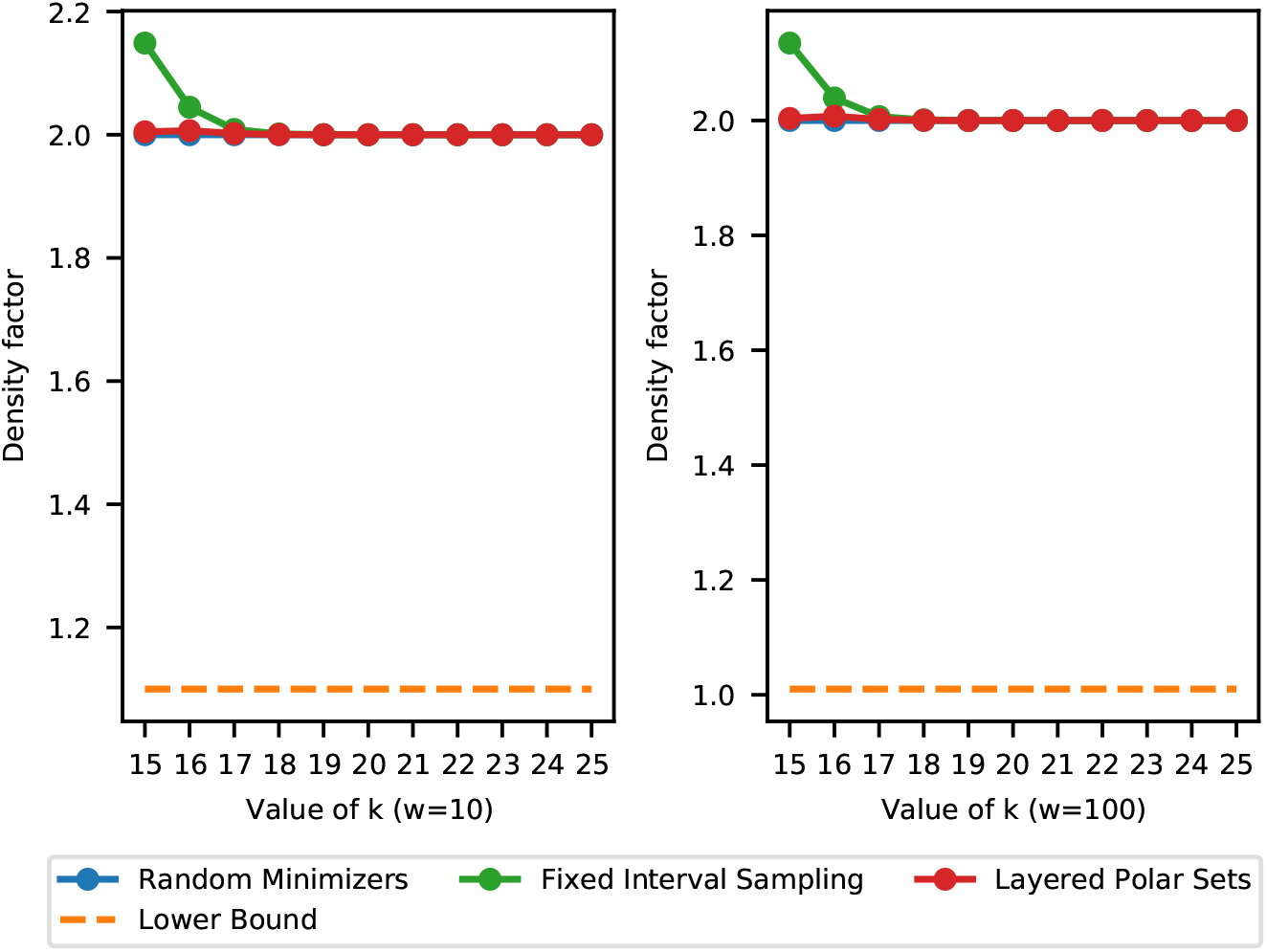
Performance of sequence-specific minimizers on random sequences (optimized on hg38) with *w* = 10 (left) and *w* = 100 (right). This is different from Figure 8: Here the specific density is measured on a unrelated random sequence.

**Figure 7:**
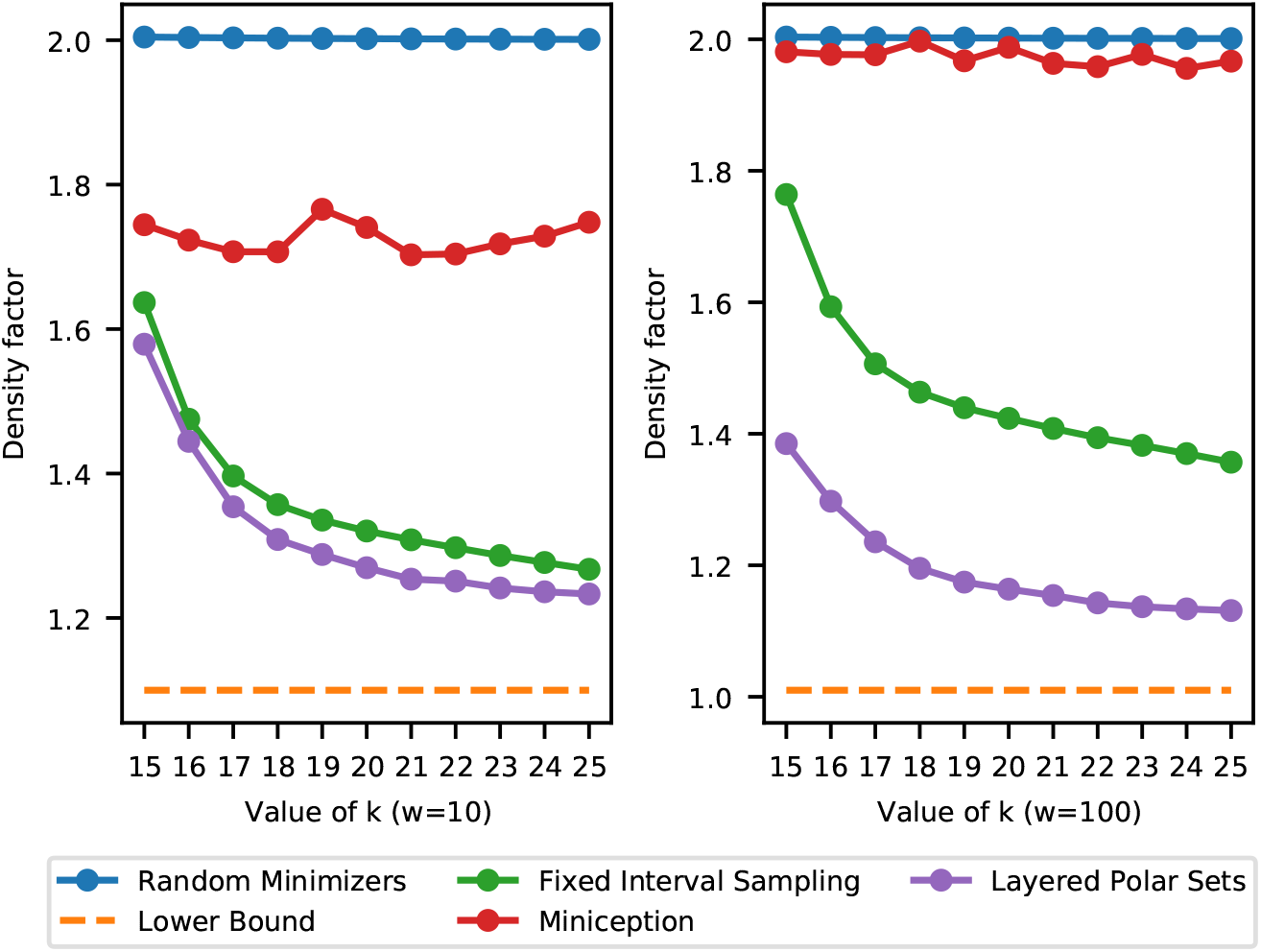
Performance of sequence-specific minimizers, optimized and tested on human chromosome 1 with *w* = 10 (left) and *w* = 100 (right).

**Figure 8:**
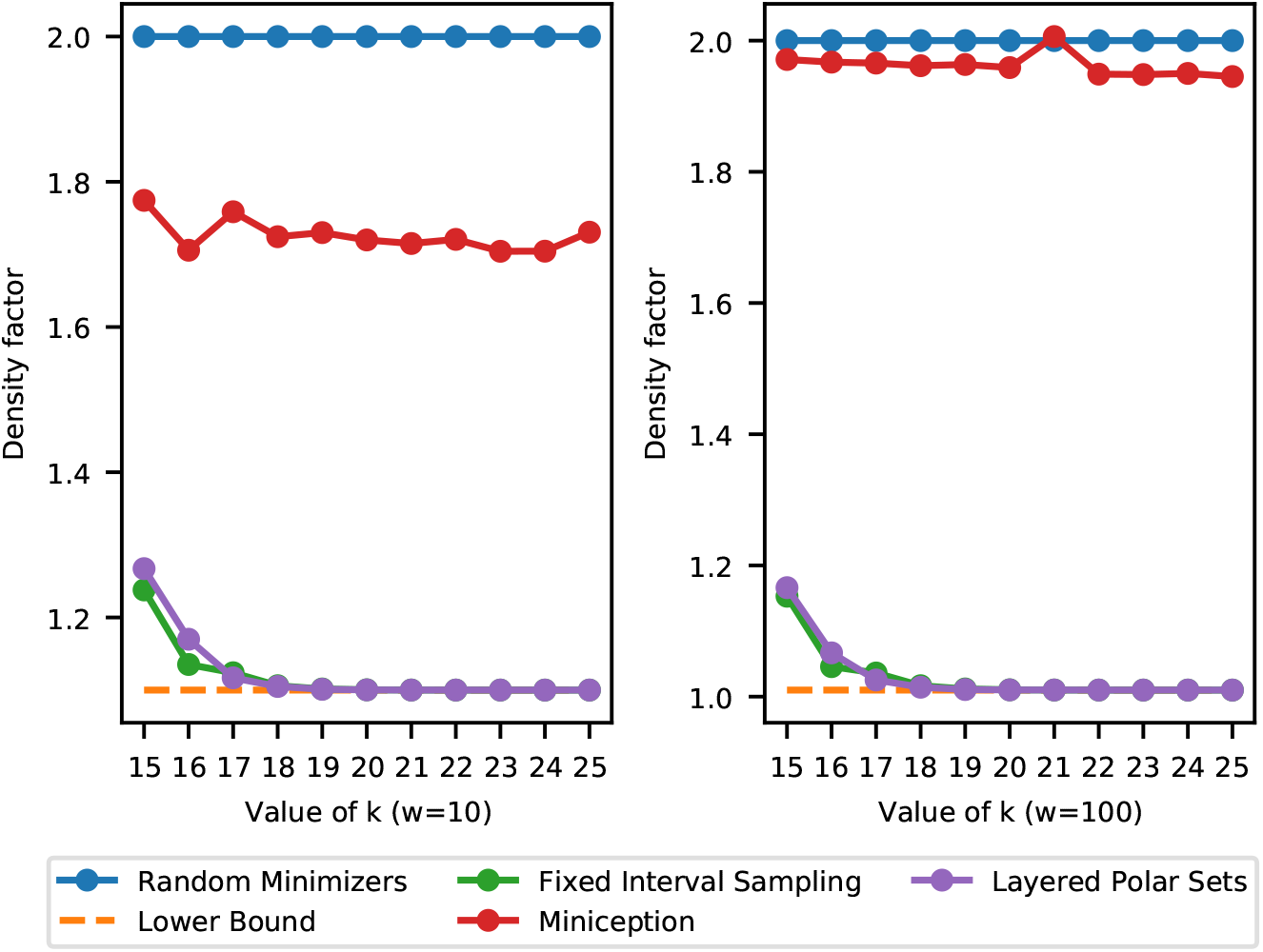
Performance of sequence-specific minimizers, optimized and tested on a 230 000 000 − long random sequence with *w* = 10 (left) and *w* = 100 (right). This is different from Figure 6: Here the specific density is measured on the same sequence the minimizers optimize on.

The reference genome is more repetitive than a purely random sequence. However, empirically the energy surplus and deficit are still small, well below 0.01 measured in density factor, implying a relative error of at most 1% when estimating specific density with link energy. Thus, when constructing efficient minimizers by (layered) polar sets, using link energy to estimate specific density is efficient and accurate. For reference, on a random sequence the average energy surplus and deficit are below 10^*−*7^ in absolute value, for the parameter range we are interested in.

### 3.2 Evaluating Polar Set Heuristics

We next evaluate our proposed algorithms for layered polar sets. We implemented the algorithm with Python3. Experiments are run in parallel and the longest ones finish within a day. The peak memory usage stands at 100 GB, which happens at the start loading the precomputed suffix array using Python pickle.

We compare our results against some other candidates:

- Random Minimizers. Achieves density factor of 2 in theory and in practice, as indicated in last section.
- Lower Bound. This corresponds to the density factor for perfect minimizers. While our theory predicts existence of perfect minimizers matching the lower bound with large value of *k*, this rarely happens with practical parameter values.
- Fixed Interval Sampling. This method uses every *w k*-mers from *S* as the set *U* to define a compatible minimizer.
- The Miniception (Zheng *et al*., 2020a), a practical algorithm that provably achieves lower density in many scenarios. The hyperparameter *k*_0_ is set to max(5, *k* − *w*) for our experiments.

We do not include existing algorithms for constructing compact universal hitting sets because these methods do not scale to values of *k >* 14. Our heuristics work the best when *k*-mers do not appear too frequently, or roughly speaking, when *σ*^*k*^ *> n* where *n* is the length of the reference sequence. This choice of parameter is common in bioinformatics analysis. With the sequence at the size of human reference genome, our heuristics work well starting at *k* = 15. Additionally, the Miniception achieves comparable performance with leading UHS-based heuristics, so its performance also serves as a viable proxy.

We consider two scenarios, first with short windows (*w* = 10) and second with long windows (*w* = 100). The results are shown in Figure 4B. Our experiments indicate that our simple heuristics yield efficient minimizers, greatly outperforming random minimizers and the Miniception, while maintaining a consistent edge over fixed interval sampling methods, in both short windows and long windows settings. The improvement is more pronounced when the windows are long. Given our layered polar set heuristics consist of multiple rounds, in Supplementary Section S5.1 we show the progression of density factors through rounds, demonstrating that the layered heuristics are particularly effective at low values of *k*. We next show that in building sequence-specific minimizers using layered anchor sets, we do not sacrifice their performance in the general case measured by (expected) density. In Supplementary Section S5.2, we sketch a random sequence using the sequence-specific minimizers we built for hg38. As expected, the performance closely matches that of a random minimizer.

## 4 Discussion

### 4.1 Limits and Future of Polar Sets

While the concept of polar sets is interesting and leads to improvements in state-of-the-art sequence-specific minimizer design, we should acknowledge its limitations. First, it cannot be used in designing non-sequence-specific minimizers when *w > k*. Arguably, this means the method is more tailored for sequence-specific minimizers. See Supplementary Section S4 for proof and more discussion on non-sequence-specific polar sets.

Our experimental results show that the performance of minimizers based on polar sets greatly improves as *k* grows. When each *k*-mer appears many times in the reference sequence, it becomes hard to select many *k*-mers without violating the polar set condition. For comparison, in Supplementary Section S5.3 we show the results when we apply the heuristics to human chromosome 1 sequence only, which is about 1/10 as long as the whole human reference genome. Improvements across the board for the heuristic algorithms and the fixed interval sampling methods are observed. The repetitiveness of human reference genome also means much more difficult optimization of specific density. In Supplementary Section S5.4, we show the results when we apply the heuristics to build sequence-specific minimizers on a random sequence that are as long as the chr1 sequence. It is significantly easier to reach the theoretical minimum specific density of 1*/w* in this setup compared to the previous one.

With better computing power and more efficient algorithms, it is desirable to compute an optimal polar set. Thanks to our link energy formulation, the problem of optimal polar set can be formed with integer linear programming (ILP), each *k*-mer being a binary variable. For moderately-sized reference sequences, an optimal polar set can be found. However, no such convenient formulation exists for layered polar sets, and it is an interesting question whether there is a tractable optimization problem for minimizers in general.

### 4.2 Practicality of Sketches-by-Optimization

The polar sets can be used wherever universal hitting sets are used, in most cases. Given that our heuristics for layered polar sets only produce a small number of layers, implementation of a compatible minimizer with layered polar sets is not fundamentally different from that with a universal hitting set. The fixed interval sampling method is very similar to previously proposed methods (Khiste and Ilie, 2015; Almutairy and Torng, 2018; Frith *et al*., 2020), where the sketch of a reference sequence is simply the set of *k*-mers appearing at locations divisible by *w*. Polar sets might not be able to directly replace fixed interval sampling, however it can be readily expanded into a set of seeds that covers the whole reference sequence.

These approaches are currently relatively underused, compared to more traditional approach of minimizers like lexicographical, random or slight variants of either one. A significant reason for their unpopularity is the fact that using these methods requires looking up a table of *k*-mers, be it a set of polar *k*-mers or universal hitting *k*-mers, for every *k*-mer in the query sequence. In contrast, for a random minimizer implemented using a hash function, no lookup is required during the sequence sketch generation process. Since these lookup tables are usually the result of sequence-specific optimization, we say these methods fall into the category of “sketches-by-optimization”. This contrast leads to interesting tradeoffs in efficiency. For example, using a polar-set-compatible minimizer generates a more compact sequence sketch, but might take more time at query compared to using a random minimizer, due to the time spent in loading and querying the set of polar *k*-mers.

We believe better implementation of *k*-mer lookup tables and better optimization of sequence sketches, possibly in a joint manner, will popularize sketches-by-optimization. Existing methods already take step towards this goal. Jain *et al*. (2020b) uses a compact lookup table to index frequent *k*-mers, and Liu *et al*. (2019) uses a Bloom filter to perform approximate query over fixed interval samples. Techniques like *k*-mer Bloom filters (Pellow *et al*., 2017) might also further help the performance.

### 4.3 Alternative Measurements of Efficiency

Throughout this manuscript our goal has been the optimization of specific density. Low density results in smaller sequence sketches, and for many applications this is desirable. However, depending on the way one uses the sequence sketch, alternative measurements of efficiency may be desirable (also see discussion in Edgar (2020)). For example, in *k*-mer counting, minimizers are used to place *k*-mers into buckets. In this case, the specific density is less relevant, and we are more concerned about the number of buckets, and the load balance between different buckets (Marçais *et al*., 2017; Nyström-Persson *et al*., 2020). For read mapping, smaller sequence sketches have its own advantage, while some may prefer reducing the number of matches, or reducing the false positive seed matches in general. We believe many of these objectives are correlated with each other, and we are interested in both further exploring benefits of a small sequence sketch, and optimization techniques for alternative measurements of efficiency.

## 5 Conclusion

Inspired by deficiencies with current theory and practice around sequence-specific minimizers, we propose the concept of polar sets, a new approach to construct sequence-specific minimizers with the ability to directly optimize the specific density of the resulting sequence sketch. We also propose simple and efficient heuristics for constructing (layered) polar sets, and demonstrate via experiments on the human reference genome the superior performance of minimizers constructed by our proposed heuristics. While there are still concerns around the practical utility, we believe the polar set framework will be a valuable asset in design and analysis of efficient sequence sketches.

## Supplementary Materials

### S1 A Technical Lemma on *k*-mer Repetition

Here we prove a technical lemma on repetitive occurrence of *k*-mers. Similar versions of this can be found in (Chikhi *et al*., 2015). Recall *σ* is the size of the alphabet.

**Lemma S7**. *Given a random sequence and a pair of locations i < j, the probability that the k-mer starting at i equals the k-mer starting at j is exactly σ*^*−k*^.

*Proof*. If *j* − *i* ≥ *k*, the two *k*-mers do not share bases, so given they are both random *k*-mers independent of each other, the probability is *σ*^*−k*^. Otherwise, the two *k*-mers intersect. We let *d* = *j* − *i*, and use *m*_*i*_ to denote the *k*-mer starting at location *i*. We use *s* to denote the substring from the start of *m*_*i*_ to the end of *m*_*j*_ with length *k* + *d* (or equivalently, the union of *m*_*i*_ and *m*_*j*_). If *m*_*i*_ = *m*_*j*_, the *p*^th^ character of *m*_*i*_ is equal to the *p*^th^ character of *m*_*j*_, meaning *s*_*p*_ = *s*_*p*+*d*_ for all 0 ≤ *p < k*. This further means *s* is a repeating sequence of period *d*, so *s* is uniquely determined by its first *d* characters and there are *σ*^*d*^ possible configurations of *s*. The probability a random *s* satisfies *m*_*i*_ = *m*_*j*_ is then *σ*^*d*^*/σ*^*k*+*d*^ = *σ*^*−k*^.

### S2 Universal Hitting Sets and Related Analyses

Universal hitting sets have been an important component in constructing practical minimizers. In this section, we provide a more formal and technical discussion on universal hitting sets. In Section S2.1, we formally define UHS and discuss why existing heuristics to construct UHS are not adequate for sequence-specific minimizer. In Section S2.2 and Section S2.3, we discuss the two existing methods to analyze compatible minimizers of UHSes, and show that these approaches both have issues that make them unfit for our goal. In Section S2.4 we discuss how UHSes can in fact be treated as special cases of polar sets, which may inspire new developments in this line of research.

#### S2.1 Definitions and Inelasticity of UHS

**Definition S10** (Universal Hitting Sets). *Let U be a set of k-mers. If U intersects with every w consecutive k-mers, it is a UHS over k-mers with path length w and relative size* |*U* |*/σ*^*k*^.

A decycling set is a set of *k*-mers that intersect with any sufficiently long strings. Any universal hitting sets must be a decycling set, so lower bound on the size of decycling sets applies to all universal hitting sets.

**Lemma S8** (Minimal Decycling Sets). *Any UHS over k-mers with finite path length has relative size* Ω(1*/k*).

With a universal hitting set, it is guaranteed that any compatible minimizer will only select *k*-mers within the UHS on any sequence. Currently, the most popular approach for constructing efficient minimizers is via construction of a compact universal hitting set, followed by constructing a compatible minimizer. These universal hitting sets are usually constructed by expanding from a minimal decycling set. As we have shown before (Zheng *et al*., 2020b), the Mykkeltveit MDS (Mykkeltveit, 1972), the MDS that is predominantly used as the starting point already covers all windows of length *O*(*k*^3^). Empirically, with larger value of *w* only a few *k*-mers needs to be added to satisfy the universal hitting condition. As a result, UHSes constructed for different references look like each other, and the compatible minimizers do not specialize well.

A related concern about using UHSes on specific sequences is on handling of repetitive *k*-mers. As we have discussed, repetitive *k*-mers are prevalent in human reference genome. Any universal hitting set always contains homomers like *AAA ⋯ A* as it is required to cover a sequence of all *A*s. This argument also extends to other repetitive *k*-mers. Such homomers, or repetitive *k*-mers, would then be preferred when using compatible minimizers for sequence sketching. This problem of prioritizing repetitive *k*-mers is also present in fixed interval sampling. Meanwhile, existing literature (Li and Birol, 2018; Jain *et al*., 2020b) suggests it is in fact beneficial to not select these *k*-mers for read mapping, while proposing different remedies to this issue. Our proposed methods also have the effect of avoiding repetitive *k*-mers, as these *k*-mers likely don’t pass the filtering step.

#### S2.2 Analysis via Density Upper Bound

There are two existing ways to analyze the density of compatible minimizers. The first is via the following lemma, as we have mentioned in the main text:

**Lemma S9**. *If U is a UHS over k-mers, any compatible minimizer has density at most* |*U* |*/σ*^*k*^.

This lemma is universally applicable and it does not depend on the ordering within *U*. However, this is an upper bound which becomes non-informative with *w >* 2*k* and sufficiently large *k*. Because any universal hitting set is at least as large as a minimal decycling set (Lemma S8), and a random (*w, k*) − minimizer achieves density of approximately 2*/*(*w* + 1), Lemma S10 at best tells us the compatible minimizer is no worse than a random one.

#### S2.3 Analysis via Probability of Single UHS Contexts

There is a second approach to analysis of compatible minimizers from universal hitting sets (Marçais *et al*., 2017). The key lemma reads as follows (slightly rephrased):

**Lemma**. *If U is a UHS over k-mers, let SP* (*U*) *be the probability that a context contains only one element in U*. *Under certain assumptions, the expected density of a random minimizer compatible with U is* 2(1 − *SP* (*U*))*/*(*w* + 1).

We now show this lemma depends on assumptions that highly depends on the structure of *U*.

We start with some notations, slightly different from the original paper. Fix a context, let *m*_*i*_ denote the *i*^th^ *k*-mer in the context. We also let *z*_*i*_ = **1**(*m*_*i*_ ∈ *U*), let *H* denote the event that the context is charged, and let 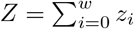 Let *C*(*n, k*) be the binomial coefficients. The proof involves the following equation (we only list the first term -there are four analogous terms):

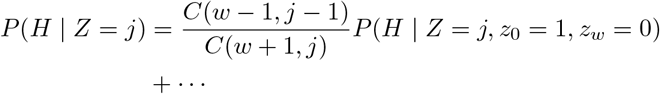

which involves a counting argument: Given 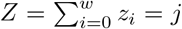, there are *C*(*w* + 1, *j*) different configurations of *z*, and *C*(*w* − 1, *j* − 1) of them satisfies *z*_0_ = 1 and *z*_*w*_ = 0. However, by invoking this counting argument, it is implicitly assumed that every configuration satisfying *?z*_*i*_ = *j* happens with the same probability, as stated (again, we only keep the terms with *z*_0_ = 1 and *z*_*w*_ = 0 and hide the rest of terms):

**Assumption**. *Let P* (*z*) *be the probability of generating a random context and observing z*_*i*_ = ***1***(*m*_*i*_ ∈ *U*). *If* 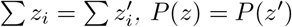.

If this is true, we also have *P* (*z* | *Z* = *j*) = 1*/C*(*w* + 1, *j*). We now recover the statement as follows:

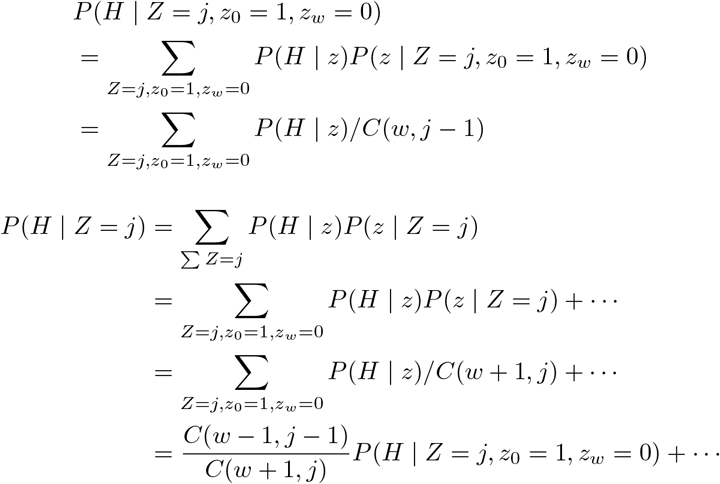

The assumption is true in expectation if the UHS itself is a random subset of Σ^*k*^, which is not the case as that set also has to satisfy the UHS condition. For a general set *U*, the probability that a *k*-mer is in *U* is highly dependent on whether the preceding intersecting *k*-mers are in *U*, and the assumption is likely not valid in most scenarios.

Finally, universal hitting sets may be constructed in a specific way to enable better analysis of compatible minimizers, as seen in (Zheng *et al*., 2020a). We do not discuss these, as they do not apply to other universal hitting sets.

#### S2.4 UHS as Improper Polar Sets

The alternative formula for link energy, as described in Section 2.3.4, allows us to define the link energy of any subset of *k*-mers, not just those satisfying the polar set condition. The main theorem for polar set still holds, but only the upper bound part. Interestingly, if we plug in a universal hitting set, we get *A*_cov_ = *n, A*_ele_ = |*U* |, *A*_seg_ = 1 and the link energy of 2*n/*(*w* + 1) |*U*| 1, where *n* is the number of *k*-mers in the reference sequence and |*U*| is the total number of times a *k*-mer in UHS appear in the reference sequence. Plugging this into the main polar set theorem, we recover the specific density upper bound |*U*| */n* for universal hitting sets, up to an error of *D*(*S*)*/n*. In this sense, universal hitting sets can be seen as a specific and extreme case of an improper polar set.

### S3 NP-Completeness of Optimal Polar Set

In this section, we show a reduction from the problem of maximal independent set to the problem of optimal polar set, with an alphabet of 3. Let *G* = (*V, E*) be the instance for maximal independent set, and without loss of generality, let |*V* |= 2^*d*^. We use Σ = {*X*, 0, 1} as the alphabet, and for the polar set instance, we let *w* = 2*d* + 1, *k* = *d* and *s* = 0. This means, we want to find subset of *d* −mers that form many links exactly 2*d* + 1 bases away, but no two *d* mers in the polar set can be fewer than 2*d* + 1 bases from each other. With *s* = 0, link energy is equivalent to number of links up to a scaling factor, so we are optimizing number of links that can be formed. We now construct the query string for polar set, which we divide into three sections.

**Disqualification Gadget**. Given an arbitrary *d* −mer *z* ∈ Σ^*d*^, we let the disqualification gadget be the following string:

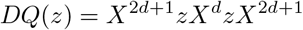

With presence of *DQ*(*z*), *z* cannot appear in the polar set, because it appears twice exactly 2*d* bases away in the disqualification gadget. The *X*^2*d*+1^ section on both ends of the gadget is to prevent *d*-mers within the gadget to form links with adjacent gadgets or sections, as *X*^*d*^ is not in the polar set.

**Disqualification Section**. We append a disqualification gadget to the query string for every *d*-mer (there are at most 3^*k*^ = *n*^1.5^ of them), **except** all *d*-mers containing only 0 and 1.

**Vertex Section**. For each vertex *v* in *G*, let *a* be its binary representation. We add *X*^2*d*+1^*aX*^*d*+1^*aX*^2*d*+1^ to the query string.

**Edge Section**. For each edge (*u, v*) in *G*, let *a, b* be the binary representation of the two ends. We add *X*^2*d*+1^*aX*^*d*^*bX*^2*d*+1^ to the query string.

The final query string is formed by the concatenation of three sections.

**Theorem S2**. *The maximal independent set can be solved by solving the optimal polar set of aforementioned query string*.

*Proof*. We claim any polar set of the query string corresponds to an independent set *V* ^*′*^ of *G*, with |*V* ^*′*^| links. All *d*-mer in the polar set are those representing vertexes in *G*, as other *d*-mers (those containing *X*) cannot appear due to the disqualification section. For each *d*-mer in the polar set, we get one link from the vertex section of the query string. If (*u, v*) ϵ *E*, the two *d*-mers representing *u* and *v* cannot be selected into the polar set at the same time, because in the edge section these two *d*-mers are apart by exactly 2*d* bases, violating polar set condition. On the other hand, all independent sets of *G* can be represented by a polar set, with total links |*V* ^*′*^| using the same argument.

We conclude that the optimal polar set of the query string is representation of a maximal independent set of *G*, which proves the statement.

This reduction also implies hardness of approximately solving optimal polar sets.

### S4 On Non-Sequence-Specific Polar Sets

For the sake of simplicity, in this section we only discuss polar sets with *s* = 0. The discussion about *s >* 0 is highly similar. As we have discussed, the density of a minimizer is the expected specific density over a random sequence. Equivalently, it equals the specific density on the de Bruijn sequence of order at least *w* + *k*. Therefore, one may construct polar sets on the de Bruijn sequence of sufficient order, to build non-sequence-specific minimizers. However, this is impossible with long windows:

**Lemma S10**. *No non-trivial polar set exists when w > k and S is the de Bruijn sequence of order w* + *k. Proof*. We simply show no *k*-mers can be in the set. For every *k*-mer *m*, the sequence *mm* exists within *S*, because *S* is the de Bruijn sequence of order at least 2*k*. Picking *m* violates the condition for polar set because it appears twice with *k < w* bases apart in *S*.

Polar sets exist on de Bruijn sequences of order *w* + *k*, when *w* ≤ *k*. With *w* = *k*, these polar sets become *non-overlapping k-mers* (Levenshtein, 1970), that is, the set of *k*-mers where no proper prefix of a *k*-mer equals a proper suffix of another *k*-mer. The problem of finding large set of non-overlapping *k*-mers is hard in general, although constructive algorithms exist (Blackburn, 2015) for constant factor approximation. With *w < k* we obtain *minimally-overlapping k-mers*, a concept that has also been studied in other contexts (Frith *et al*., 2020). We believe the concept of non-sequence-specific polar sets is of both practical and theoretical interest.

### S5 Supplementary Experiments and Figures

#### S5.1 Density Factor of Layered Polar Sets by Round

To show that our proposed layered anchor set heuristics is useful, in Figure 5 we plot the density factor after each round of optimization on the human reference genome hg38. All algorithms are run for a total of 7 rounds, with last two being monotonic rounds. We select 7 to ensure the resulting sets are not too complicated and can be computed in reasonable amount of time. With more rounds, many of the results can be further improved.

#### S5.2 Viability of Sequence-Specific Minimizers on Non-target sequences

To validate that optimization of sequence-specific density does not come at the cost of higher (non-sequence-specific) density, we generate the sequence-specific minimizers for hg38 reference genome, then apply these minimizers on a random sequence. Figure 6 shows the results. We expect these to perform close to random minimizers when *σ*^*k*^ ≫*N* where *N* is the length of the reference sequence. In these cases, most *k*-mers in a random sequence is not seen in the reference sequence, and optimized sequence-specific minimizers behave just like random minimizers in most cases. The performance for the Miniception is almost identical to that in hg38, and is not shown in this plot. The layered polar sets is also arguably more robust at lower values of *k*, as its density stays close to that of a random minimizer.

#### S5.3 Experiments on Human Chromosome 1

To show the effect of reference sequence length on the performance of sequence-specific minimizers, in Figure 7 we show the performance plot when we build sequence-specific minimizers for chr1 only. The human chromosome 1 sequence is around 10% of the whole hg38 sequence, and consistent with our theory, the time and memory spent to run these experiments on chr1 are around 10% of that for hg38 ones.

#### S5.4 Building Sequence-Specific Minimizers on Random Sequences

To further show that human reference genome is highly repetitive and construction of efficient sequence-specific minimizers is hard in such setup, we run the algorithms to generate sequence-specific minimizers on a random sequence of length 230 000 000, similar to that of chromosome 1. Figure 8 shows the performance of layered polar sets and fixed interval sampling method. Compared with Figure 7, we observe it is much easier to build efficient minimizers on a random sequence, and to match the theoretical lower bound, even given the reference sequences has similar length.

## References

Almutairy, M. and Torng, E. (2018). Comparing fixed sampling with minimizer sampling when using k-mer indexes to find maximal exact matches. PLOS ONE, 13(2), e0189960.

Blackburn, S. R. (2015). Non-overlapping codes. IEEE Transactions on Information Theory, 61(9), 4890–4894.

Chikhi, R., Limasset, A., and Medvedev, P. (2015). Compacting de Bruijn graphs from sequencing data quickly and in low memory. Bioinformatics, 32(12), i201–i208.

DeBlasio, D., Gbosibo, F., Kingsford, C., and Marçais, G. (2019). Practical universal k-mer sets for minimizer schemes. In Proceedings of the 10th ACM International Conference on Bioinformatics, Computational Biology and Health Informatics, BCB ‘19, pages 167–176, New York, NY, USA. ACM.

Deorowicz, S., Kokot, M., Grabowski, S., and Debudaj-Grabysz, A. (2015). KMC 2: Fast and resource-frugal k-mer counting. Bioinformatics, 31(10), 1569–1576.

Edgar, R. C. (2020). Syncmers are more sensitive than minimizers for selecting conserved k-mers in biological sequences. bioRxiv.

Ekim, B., Berger, B., and Orenstein, Y. (2020). A randomized parallel algorithm for efficiently finding near-optimal universal hitting sets. BioRxiv: 2020.01.17.910513.

Erbert, M., Rechner, S., and Müller-Hannemann, M. (2017). Gerbil: a fast and memory-efficient k-mer counter with GPU-support. Algorithms for Molecular Biology, 12(1), 9.

Frith, M. C., Noè, L., and Kucherov, G. (2020). Minimally-overlapping words for sequence similarity search. BioRxiv.

Jain, C., Rhie, A., Hansen, N., Koren, S., and Phillippy, A. M. (2020a). A long read mapping method for highly repetitive reference sequences. bioRxiv, page 2020.11.01.363887.

Jain, C., Rhie, A., Zhang, H., Chu, C., Walenz, B. P., Koren, S., and Phillippy, A. M. (2020b). Weighted minimizer sampling improves long read mapping. Bioinformatics, 36(Supplement 1), i111–i118.

Kempa, D. and Kociumaka, T. (2019). String synchronizing sets: sublinear-time bwt construction and optimal lce data structure. In Proceedings of the 51st Annual ACM SIGACT Symposium on Theory of Computing, pages 756–767.

Khiste, N. and Ilie, L. (2015). E-mem: efficient computation of maximal exact matches for very large genomes. Bioinformatics, 31(4), 509–514.

Levenshtein, V. I. (1970). Maximum number of words in codes without overlaps. Problemy Peredachi Informatsii, 6(4), 88–90.

Li, H. and Birol, I. (2018). Minimap2: Pairwise alignment for nucleotide sequences. Bioinformatics, 34(18), 3094–3100.

Liu, Y., Zhang, L. Y., and Li, J. (2019). Fast detection of maximal exact matches via fixed sampling of query k-mers and bloom filtering of index k-mers. Bioinformatics, 35(22), 4560–4567.

Marçais, G., Pellow, D., Bork, D., Orenstein, Y., Shamir, R., and Kingsford, C. (2017). Improving the performance of minimizers and winnowing schemes. Bioinformatics, 33(14), i110–i117.

Marçais, G., DeBlasio, D., and Kingsford, C. (2018). Asymptotically optimal minimizers schemes. Bioinformatics, 34(13), i13–i22.

Marçais, G., Solomon, B., Patro, R., and Kingsford, C. (2019). Sketching and sublinear data structures in genomics. Annual Review of Biomedical Data Science, 2(1), 93–118.

Mykkeltveit, J. (1972). A proof of Golomb’s conjecture for the de Bruijn graph. Journal of Combinatorial Theory, Series B, 13(1), 40–45.

Nyström-Persson, J. T., Keeble-Gagnère, G., and Zawad, N. (2020). Compact and evenly distributed k-mer binning for genomic sequences. bioRxiv.

Orenstein, Y., Pellow, D., Marçais, G., Shamir, R., and Kingsford, C. (2016). Compact universal k-mer hitting sets. In Algorithms in Bioinformatics, Lecture Notes in Computer Science, pages 257–268. Springer, Cham.

Pellow, D., Filippova, D., and Kingsford, C. (2017). Improving bloom filter performance on sequence data using k-mer bloom filters. Journal of Computational Biology, 24(6), 547–557.

Roberts, M., Hunt, B. R., Yorke, J. A., Bolanos, R. A., and Delcher, A. L. (2004a). A preprocessor for shotgun assembly of large genomes. Journal of Computational Biology, 11(4), 734–752.

Roberts, M., Hayes, W., Hunt, B. R., Mount, S. M., and Yorke, J. A. (2004b). Reducing storage requirements for biological sequence comparison. Bioinformatics, 20(18), 3363–3369.

Schleimer, S., Wilkerson, D. S., and Aiken, A. (2003). Winnowing: Local Algorithms for Document Fingerprinting. In Proceedings of the 2003 ACM SIGMOD International Conference on Management of Data, SIGMOD ‘03, pages 76–85. ACM.

Ye, C., Ma, Z. S., Cannon, C. H., Pop, M., and Yu, D. W. (2012). Exploiting sparseness in de novo genome assembly. BMC Bioinformatics, 13, S1.

Zheng, H., Kingsford, C., and Marçais, G. (2020a). Improved design and analysis of practical minimizers. Bioinformatics, 36(Supplement 1), i119–i127.

Zheng, H., Kingsford, C., and Marçais, G. (2020b). Lower density selection schemes via small universal hitting sets with short remaining path length. arXiv preprint arXiv:2001.06550.

